## Supplementary Figures 1-10 for "Functional Profiling of Single CRISPR/Cas9-Edited Human Long-Term Hematopoietic Stem Cells"

**Supplementary Figure Legends**

**Supplementary Figure 1: Lineage hierarchy of human hematopoietic stem cells.** Major classes of stem and progenitor cells, which are defined by cell surface markers. Long-term hematopoietic stem cells (LT-HSCs), which are defined by the expression of CD49f and other markers, constitute to only about 0.1 - 1% of bulk CD34+ hematopoietic stem and progenitor cells (HSPCs). LT-HSCs give rise to various types of progenitor cells including short-term hematopoietic stem cells (ST-HSCs), multi-lymphoid progenitor cells (MLPs), common myeloid progenitor cells (CMPs), granulocyte/macrophage progenitor cells (GMPs) and myelo–erythroid progenitor cells (MEPs). MEPs terminally differentiate into erythrocytes and megakaryocytes.

**Supplementary Figure 2: Flow cytometry sorting scheme.** Lineage depleted neonatal cord blood was utilized to sort LT-HSCs as $CD45^{+}CD34^{+}CD38^{-}CD45RA^{-}CD90^{+}CD49f^{+}$, ST-HSCs as $CD45^{+}CD34^{+}CD38^{-}CD45RA^{-}CD90^{-}CD49f^{-}$ and MEPs as $CD45^{+}CD34^{+}CD38^{+}CD10/19^{-}CD7^{-}CD45RA^{-}FLT3^{-}$.

**Supplementary Figure 3: CRISPR/Cas9-mediated isoform assignment to GATA1-Short and –Long**. **a** Gel electrophoresis analysis of single cell-derived colonies that were CRISPR/Cas9 edited with control gRNAs. Only homozygous knock-out colonies were utilized in the analysis. **b** Gel electrophoresis analysis of single cell-derived colonies that were CRISPR/Cas9 edited for GATA1-Short. GATA1 is located on the X-chromosome and only male cord blood samples were utilized. **c** Sanger sequencing analysis of single cell-derived colonies that were CRISPR/Cas9 edited for GATA1-Long. The alternative start site was mutated from ATG to CTC and the PAM sequence mutated from GGG to GGC to avoid repeated cutting by the gRNA.

**Supplementary Figure 4: Single cell *in vitro* differentiation assay. a** Off-target cleavage efficiency of genomic loci that were similar in sequence to the gRNA target sequence ($n$ = 5-10 single cell-derived colonies per genomic loci). **b** Percentage of CRISPR/Cas9 efficiency as determined by single cell-derived colonies that were positive for STAG2 knock-out (only frame shift mutations). STAG2 is located on the X-chromosome and only male cord blood samples were utilized ($n$ = 3 experiments with independent cord blood pools and >50 colonies per experiment, error bars represent standard deviations). **c** Sanger sequencing analysis of single cell-derived colonies that were CRISPR/Cas9 edited for STAG2 knock-out. Example of a frame shift mutation is depicted.

**Supplementary Figure 5: Setup of near-clonal xenotransplantation in NSG mice. a** Limiting dilution analysis (LDA) of LT-HSCs transplanted NSG mice at varying doses for 20 weeks revealed a stem cell frequency of ~1/100 ($n$ = 1 experiment). **b** Percentage of control and GATA1-Short edited LT-HSCs injected NSG mice with engraftment of >5% in the RF (based on human $CD45^+$ expression, $n$ = 3 experiments with independent cord blood pools, error bars represent standard deviations, which is also the case for **c**). **c** Percentage of control and GATA1-Short edited LT-HSCs injected NSG mice with high CRISPR/Cas9 knock-out efficiency (>90% based on PCR and Sanger sequencing). **d** Analysis of chromatograms from cells of the RF of control and GATA1-Short edited LT-HSCs injected NSG mice. Percentage of aberrant sequences at the gRNA cut site were utilized to assess CRISPR/Cas9 efficiency. **e** Examples of chromatograms from cells of the RF of control and GATA1-Short edited LT-HSCs injected NSG mice. Note the single clonal engraftment of a GATA1-Short edited LT-HSCs injected NSG mice based on the single chromatogram trace (second from top).

**Supplementary Figure 6: Methylcellulose colony formation from LT-HSCs transplanted mice.** Sanger sequencing of individual colonies from methylcellulose colony formation and their frequency. No wildtype GATA1 colonies were detected ($n$ = 5 mice).

**Supplementary Figure 7: Analysis of near-clonal xenotransplantation in NSG mice. a** Percentage of $CD45^{+}CD33^{+}$ myeloid cells in control and GATA1-Short edited LT-HSCs injected NSG mice in RF and BM ($n$ = 3 experiments with independent cord blood pools, error bars represent standard deviations, which is also the case for **b-i**, unpaired t-test $P < 0.05$ for RF GATA1-Short versus RF control). **b** Percentage of $CD45^{-}GlyA^{+}$ erythroid cells in control and GATA1-Short edited LT-HSCs injected NSG mice in the RF and BM. **c** Percentage of $CD45^{+}CD3^{+}$ T-cells in control and GATA1-Short edited LT-HSCs injected NSG mice in the RF and BM. **d** Percent distribution of all cell lineages in control and GATA1-Short edited LT-HSCs injected NSG mice in RF and BM. **e** Absolute cell numbers of $CD45^+CD19^+$ lymphoid cells in control and GATA1-Short edited LT-HSCs injected NSG mice in RF and BM (unpaired t-test $P$ < 0.01 for RF GATA1-Short versus RF control). **f** Absolute cell numbers of $CD45^+CD33^+$ myeloid cells in control and GATA1-Short edited LT-HSCs injected NSG mice in RF and BM. **g** Absolute cell numbers of $CD45^-CD41^+$ megakaryocytes in control and GATA1-Short edited LT-HSCs injected NSG mice in RF and BM (unpaired t-test $P$ < 0.005 for RF GATA1-Short versus RF control). **h** Absolute cell numbers of $CD45^-$ $GlyA^+$ erythroid cells in control and GATA1-Short edited LT-HSCs injected NSG mice in RF and BM. **i** Absolute cell numbers of $CD45^+CD3^+$ T-cells in control and GATA1-Short edited LT-HSCs injected NSG mice in RF and BM.

**Supplementary Figure 8: Setup of near clonal xenotransplantation in NSGW41 mice.** **a** Limiting dilution analysis (LDA) of LT-HSCs transplanted NSGW41 mice at varying doses for 12 weeks revealed a stem cell frequency of ~1/175 ($n$ = 1 experiment). **b** Percentage of control and GATA1-Short edited LT-HSCs injected NSGW41 mice with engraftment of >5% in the RF (based on human $CD45^+$ expression, $n$ = 3 experiments with independent cord blood pools, error bars represent standard deviations, which is also the case for **c**). **c** Percentage of control and GATA1-Short edited LT-HSCs injected NSGW41 mice with high CRISPR/Cas9 knock-out efficiency (>90% based on PCR and Sanger sequencing).

**Supplementary Figure 9: Analysis of near clonal xenotransplantation in NSGW41 mice.** **a** Percentage of $CD45^+CD19^+$ lymphoid cells in control and GATA1-

Short edited LT-HSCs injected NSGW41 mice in RF and BM ($n$ = 3 experiments with independent cord blood pools, error bars represent standard deviations, which is also the case for **b-g**). **b** Percentage of $CD45^+CD33^+$ myeloid cells in control and GATA1-Short edited LT-HSCs injected NSGW41 mice in RF and BM. **c** Percent distribution of all cell lineages in control and GATA1-Short edited LT-HSCs injected NSGW41 mice in RF and BM. **d** Absolute cell numbers of $CD45^+CD19^+$ lymphoid cells in control and GATA1-Short edited LT-HSCs injected NSGW41 mice in RF and BM. **e** Absolute cell numbers of $CD45^+CD33^+$ myeloid cells in control and GATA1-Short edited LT-HSCs injected NSGW41 mice in RF and BM. **f** Absolute cell numbers of $CD45^-CD41^+$ megakaryocytes in control and GATA1-Short edited LT-HSCs injected NSGW41 mice in RF and BM (unpaired t-test $P < 0.005$ for GATA1-Short versus control for both RF and BM). **g** Absolute cell numbers of $CD45^-GlyA^+$ erythroid cells in control and GATA1-Short edited LT-HSCs injected NSGW41 mice in RF and BM (unpaired t-test $P < 0.005$ for GATA1-Short versus control in both RF and BM).

**Supplementary Figure 10: Full length gel pictures and western assay. a** Gel electrophoresis picture from Supplementary Fig. 3a. **b** Gel electrophoresis picture from Supplementary Fig. 3b. **c** Western assay image from Fig. 2c.

Wagenblast et al. Supplementary Figure 1

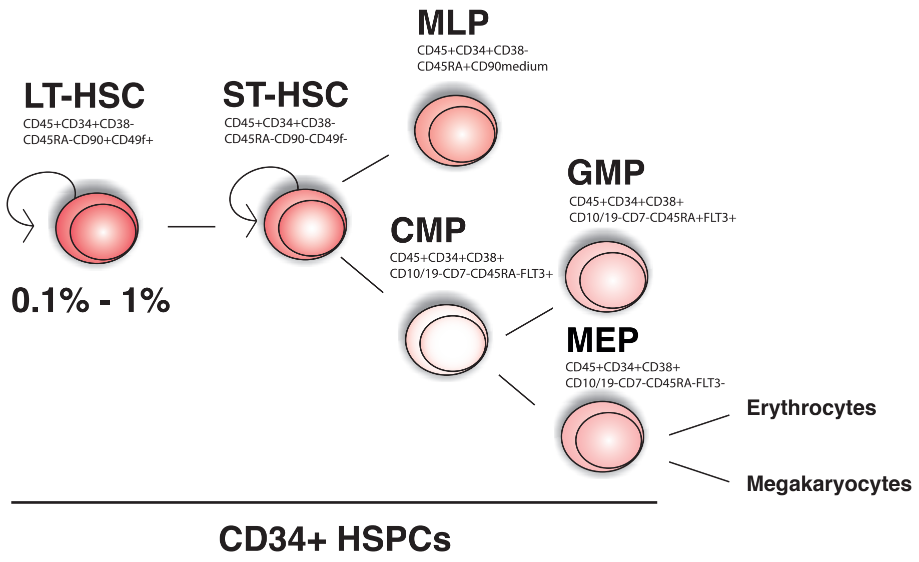

Wagenblast et al. Supplementary Figure 2

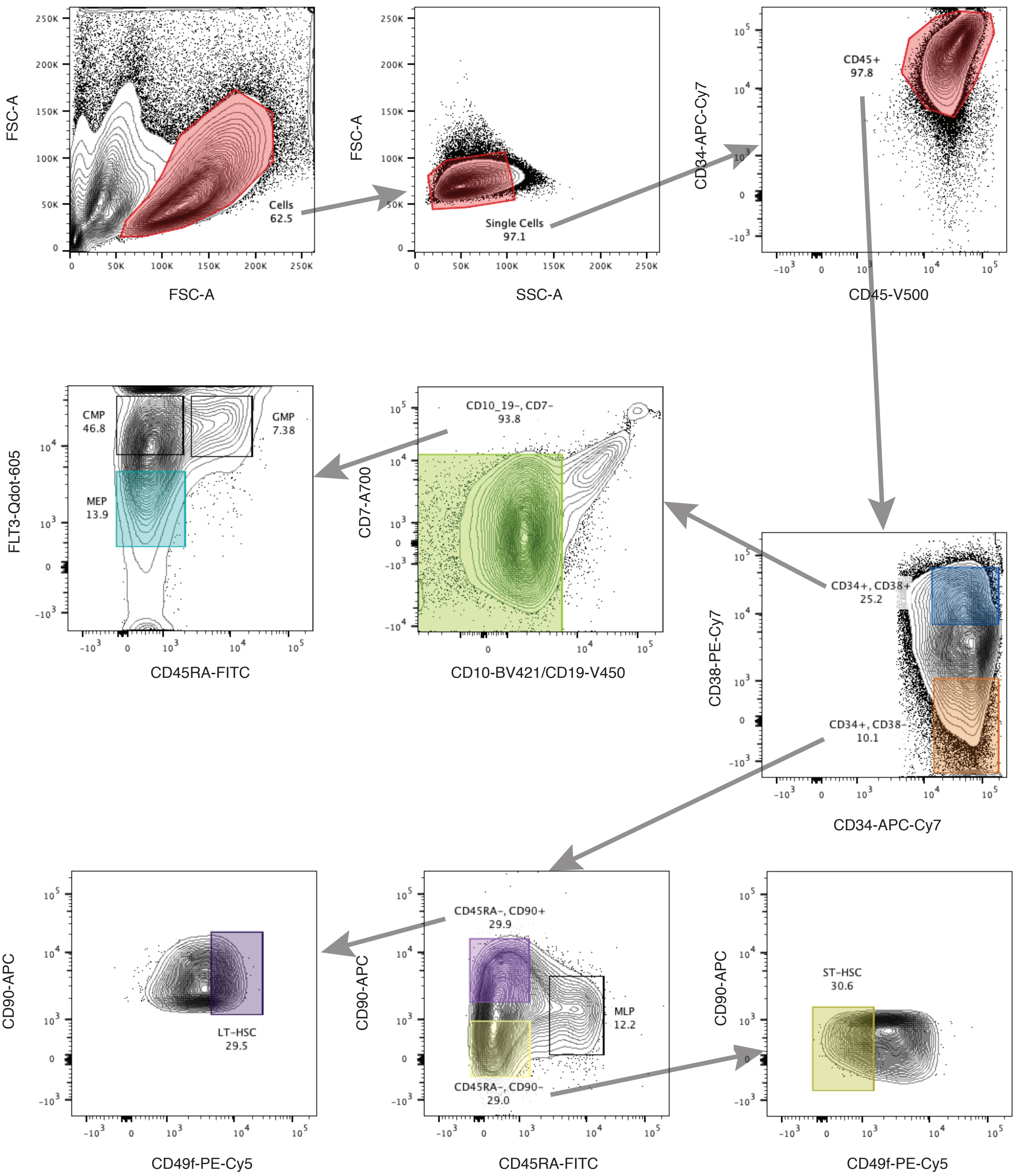

Wagenblast et al. Supplementary Figure 3

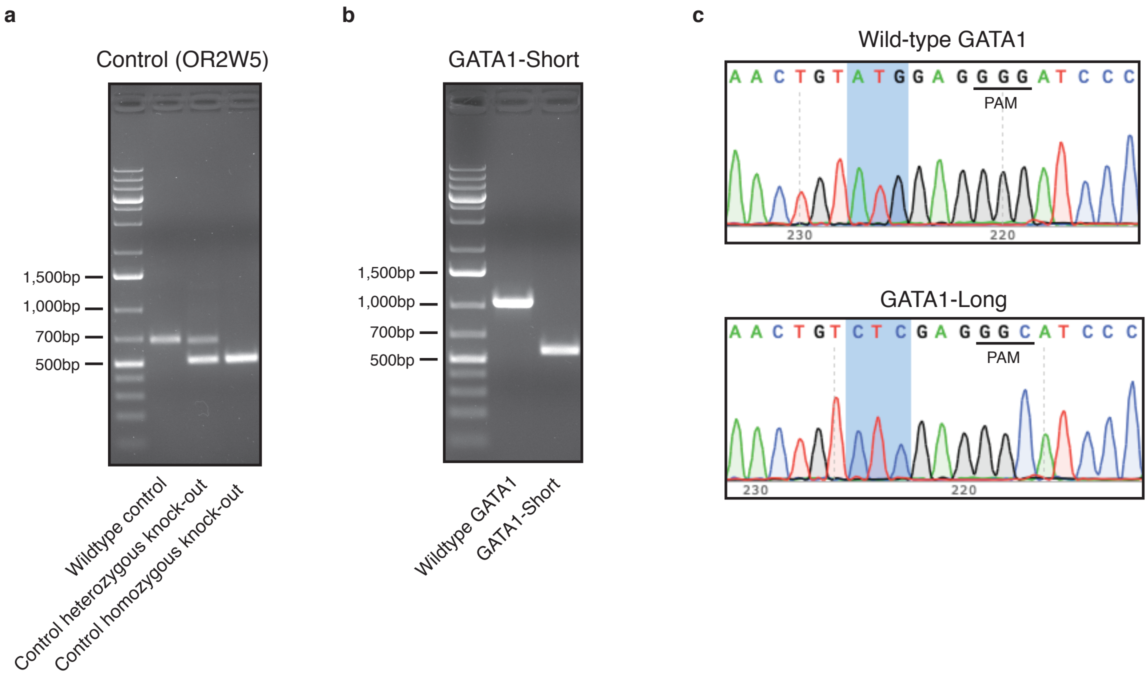

Wagenblast et al. Supplementary Figure 4

**a**

| gRNA | Colony | Chromosome location | % Cleavage efficiency (measured by TIDE) |
|---|---|---|---|
| GATA1-Short gRNA-1 | LT-HSC | chr10_20221643 | No cleavage |
| GATA1-Short gRNA-1 | ST-HSC | chr10_20221643 | No cleavage |
| GATA1-Short gRNA-1 | MEP | chr10_20221643 | No cleavage |
| GATA1-Short gRNA-1 | LT-HSC | chr18_8376641 | No cleavage |
| GATA1-Short gRNA-1 | ST-HSC | chr18_8376641 | No cleavage |
| GATA1-Short gRNA-1 | MEP | chr18_8376641 | No cleavage |
| GATA1-Short gRNA-2 | LT-HSC | chr8_107174119 | No cleavage |
| GATA1-Short gRNA-2 | ST-HSC | chr8_107174119 | No cleavage |
| GATA1-Short gRNA-2 | MEP | chr8_107174119 | No cleavage |
| GATA1-Short gRNA-2 | LT-HSC | chr12_2085541 | No cleavage |
| GATA1-Short gRNA-2 | ST-HSC | chr12_2085541 | No cleavage |
| GATA1-Short gRNA-2 | MEP | chr12_2085541 | No cleavage |
| GATA1-Short gRNA-2 | LT-HSC | chr3_127784219 | No cleavage |
| GATA1-Short gRNA-2 | ST-HSC | chr3_127784219 | No cleavage |
| GATA1-Short gRNA-2 | MEP | chr3_127784219 | No cleavage |
| GATA1-Short gRNA-2 | LT-HSC | chr7_17679912 | No cleavage |
| GATA1-Short gRNA-2 | ST-HSC | chr7_17679912 | No cleavage |
| GATA1-Short gRNA-2 | MEP | chr7_17679912 | No cleavage |
| GATA1-Long gRNA-1 | LT-HSC | chr12_20017977 | No cleavage |
| GATA1-Long gRNA-1 | ST-HSC | chr12_20017977 | No cleavage |
| GATA1-Long gRNA-1 | MEP | chr12_20017977 | No cleavage |
| GATA1-Long gRNA-1 | LT-HSC | chr12_94844478 | No cleavage |
| GATA1-Long gRNA-1 | ST-HSC | chr12_94844478 | No cleavage |
| GATA1-Long gRNA-1 | MEP | chr12_94844478 | No cleavage |
| GATA1-Long gRNA-1 | LT-HSC | chr2_15662510 | No cleavage |
| GATA1-Long gRNA-1 | ST-HSC | chr2_15662510 | No cleavage |
| GATA1-Long gRNA-1 | MEP | chr2_15662510 | No cleavage |
| GATA1-Long gRNA-1 | LT-HSC | chr9_76619884 | No cleavage |
| GATA1-Long gRNA-1 | ST-HSC | chr9_76619884 | No cleavage |
| GATA1-Long gRNA-1 | MEP | chr9_76619884 | No cleavage |
| Control gRNA-1 | LT-HSC | chr2_30448683 | No cleavage |
| Control gRNA-1 | ST-HSC | chr2_30448683 | No cleavage |
| Control gRNA-1 | MEP | chr2_30448683 | No cleavage |
| Control gRNA-1 | LT-HSC | chr6_169886852 | No cleavage |
| Control gRNA-1 | ST-HSC | chr6_169886852 | No cleavage |
| Control gRNA-1 | MEP | chr6_169886852 | No cleavage |
| Control gRNA-1 | LT-HSC | chr20_52425093 | No cleavage |
| Control gRNA-1 | ST-HSC | chr20_52425093 | No cleavage |
| Control gRNA-1 | MEP | chr20_52425093 | No cleavage |
| Control gRNA-2 | LT-HSC | chr15_54082949 | No cleavage |
| Control gRNA-2 | ST-HSC | chr15_54082949 | No cleavage |
| Control gRNA-2 | MEP | chr15_54082949 | No cleavage |

**b**

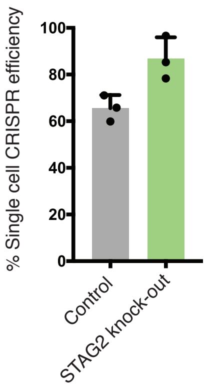

**c**

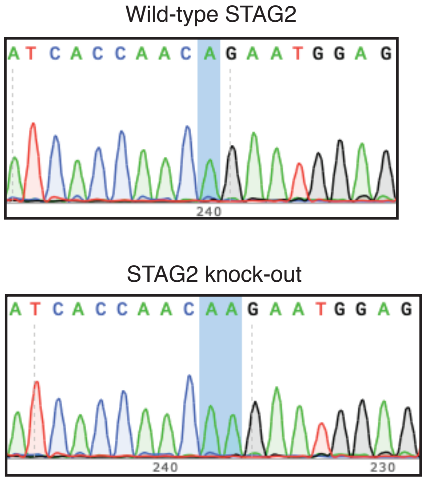

Wagenblast et al. Supplementary Figure 5

**a**

| NSG | LT-HSC dose | Transplantation engrafted/injected | Stem cell frequency |
|---|---|---|---|
| Control | 25 | 2/10 | 1/104 (Range: 65.5 - 164) |
| | 50 | 4/10 | |
| | 100 | 7/10 | |
| | 200 | 8/10 | |

**b**

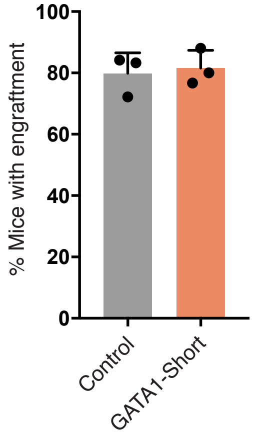

**c**

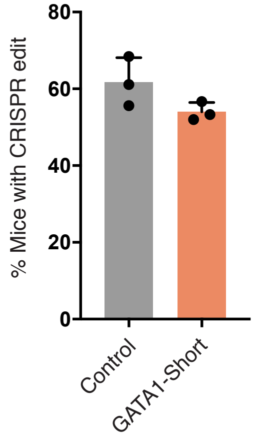

**d**

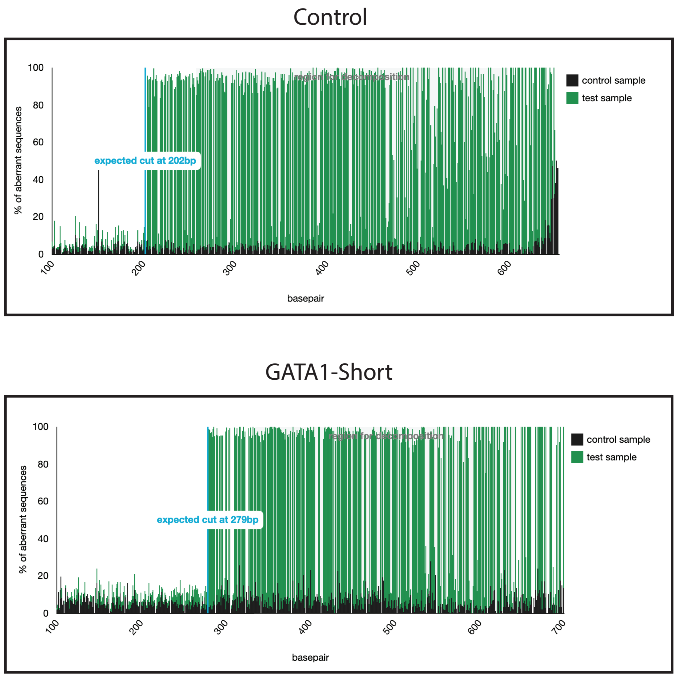

**e**

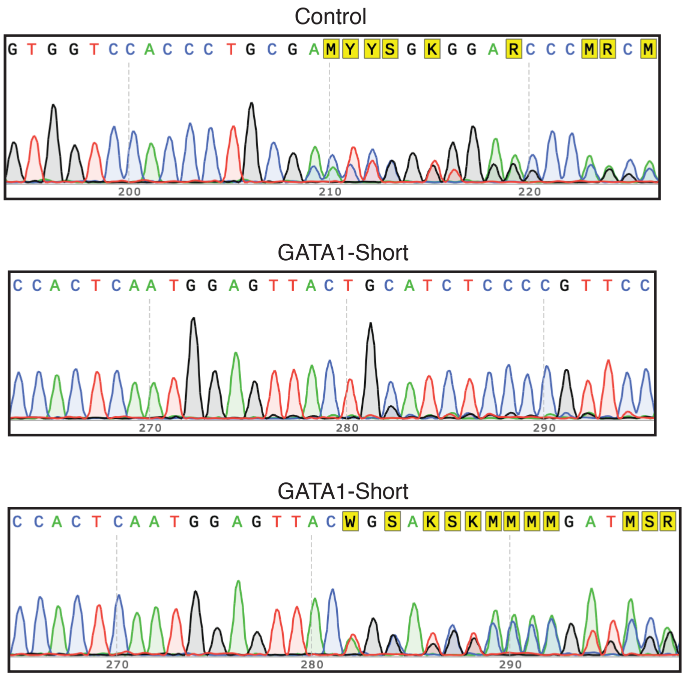

| Mouse | Sequence of methylcellulose colonies (gRNA-1 to -2 junction) | Frequency |
|---|---|---|
| 1 | 5'-TGGAACGGGGAGATGCAGTAACTCCATTGAGTGG-3' | 24/24 |
| 2 | 5'-TGGAACGGGGAGATGTAACTCCATTGAGTGG-3' | 20/20 |
| 3 | 5'-TGGAACGGGGAGATGCGTAACTCCATTGAGTGG-3' | 22/22 |
| 4 | 5'-TGGAACGGGGAGATGCAGTAACTCCATTGAGTGG-3' | 17/22 |
| | 5'-TGGAACGGGGAGATGCAACTCCATTGAGTGG-3' | 5/22 |
| 5 | 5'-TGGAACGGGGAGGTAACTCCATTGAGTGG-3' | 14/22 |
| | 5'-TGGAACGGGGAGATGCCACTCCATTGAGTGG-3' | 8/22 |

Wagenblast et al. Supplementary Figure 7

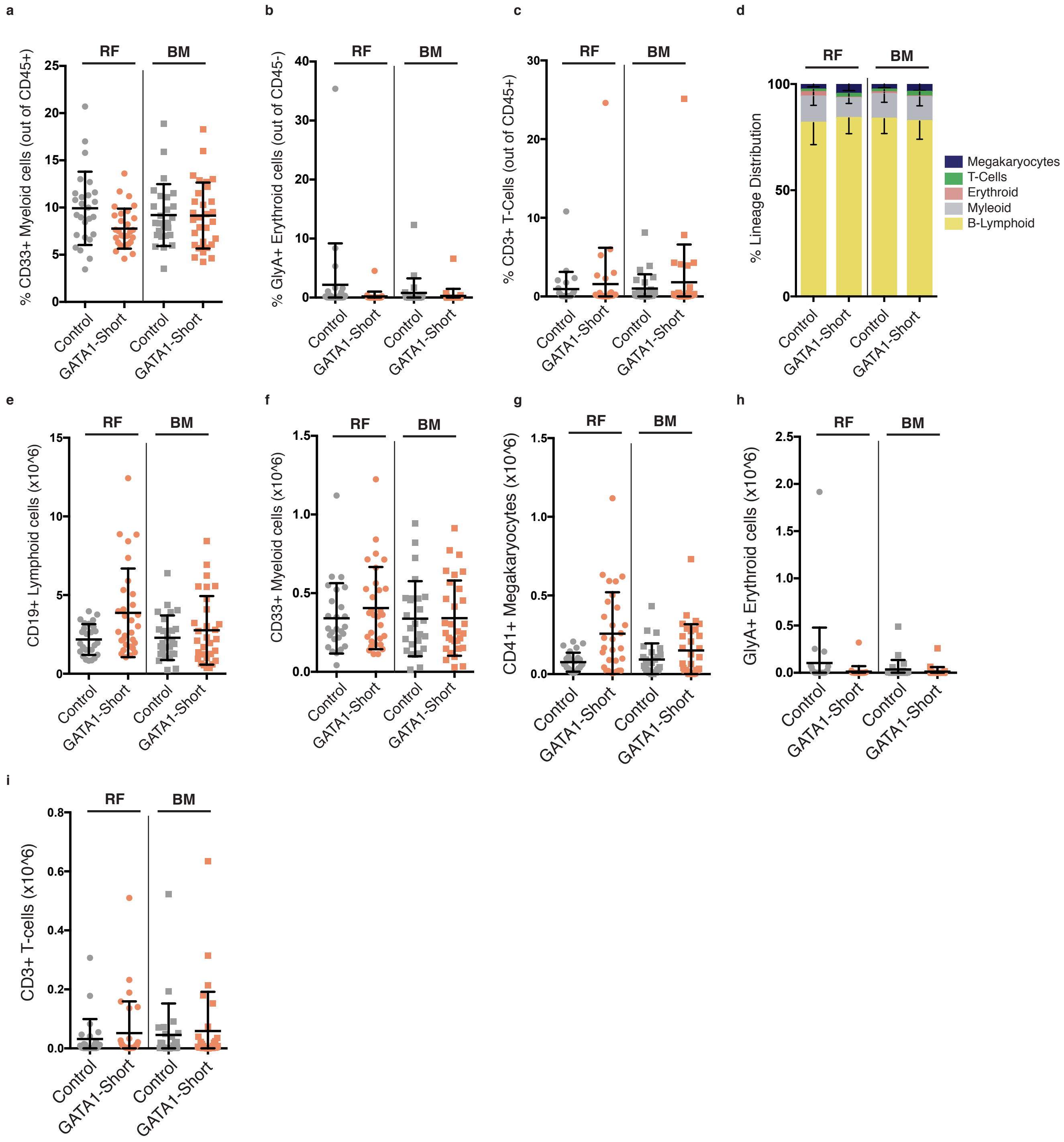

Wagenblast et al. Supplementary Figure 8

a

| NSGW41 | LT-HSC dose | Transplantation engrafted/injected | Stem cell frequency |
|---|---|---|---|
| Control | 25 | 2/10 | 1/175 |
| | 50 | 2/10 | |
| | 100 | 4/10 | |
| | 200 | 7/10 | (Range: 104 - 295) |

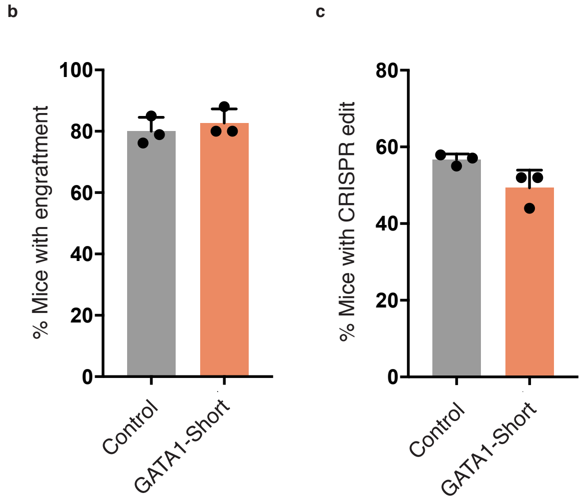

Wagenblast et al. Supplementary Figure 9

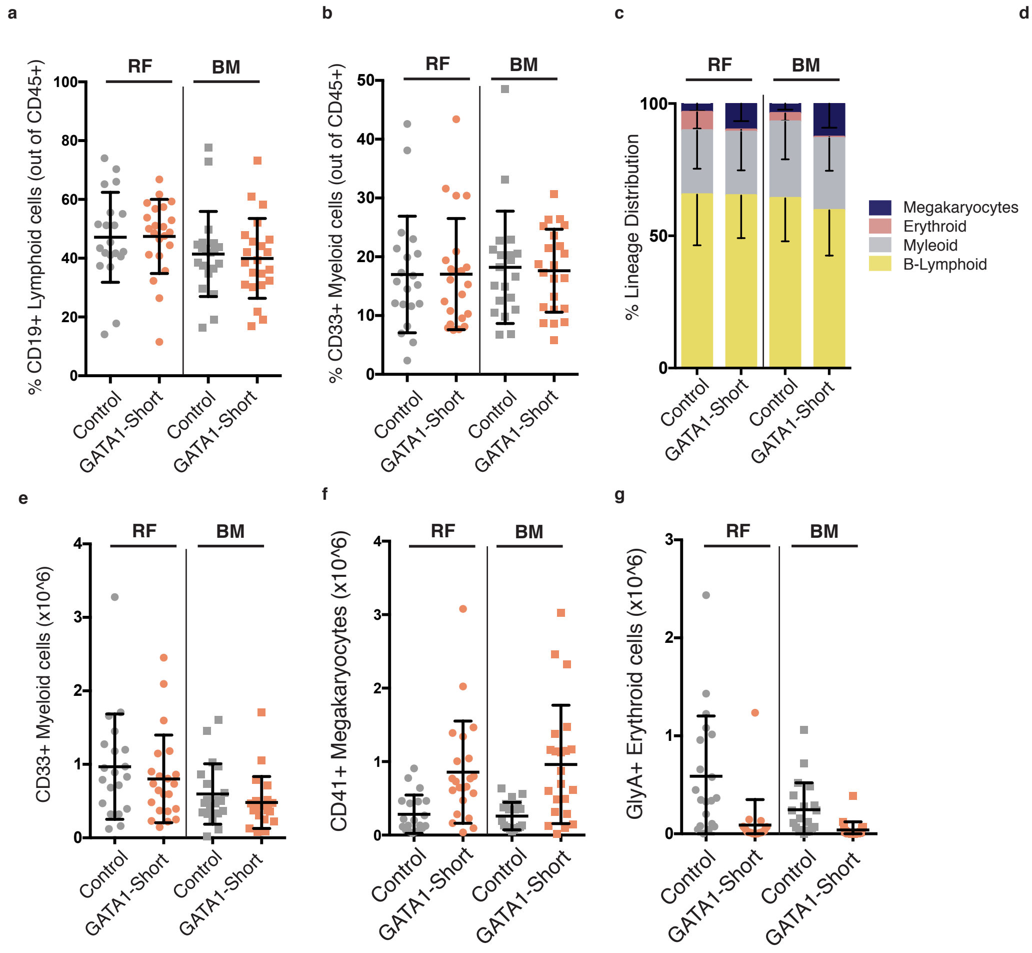

Wagenblast et al. Supplementary Figure 10

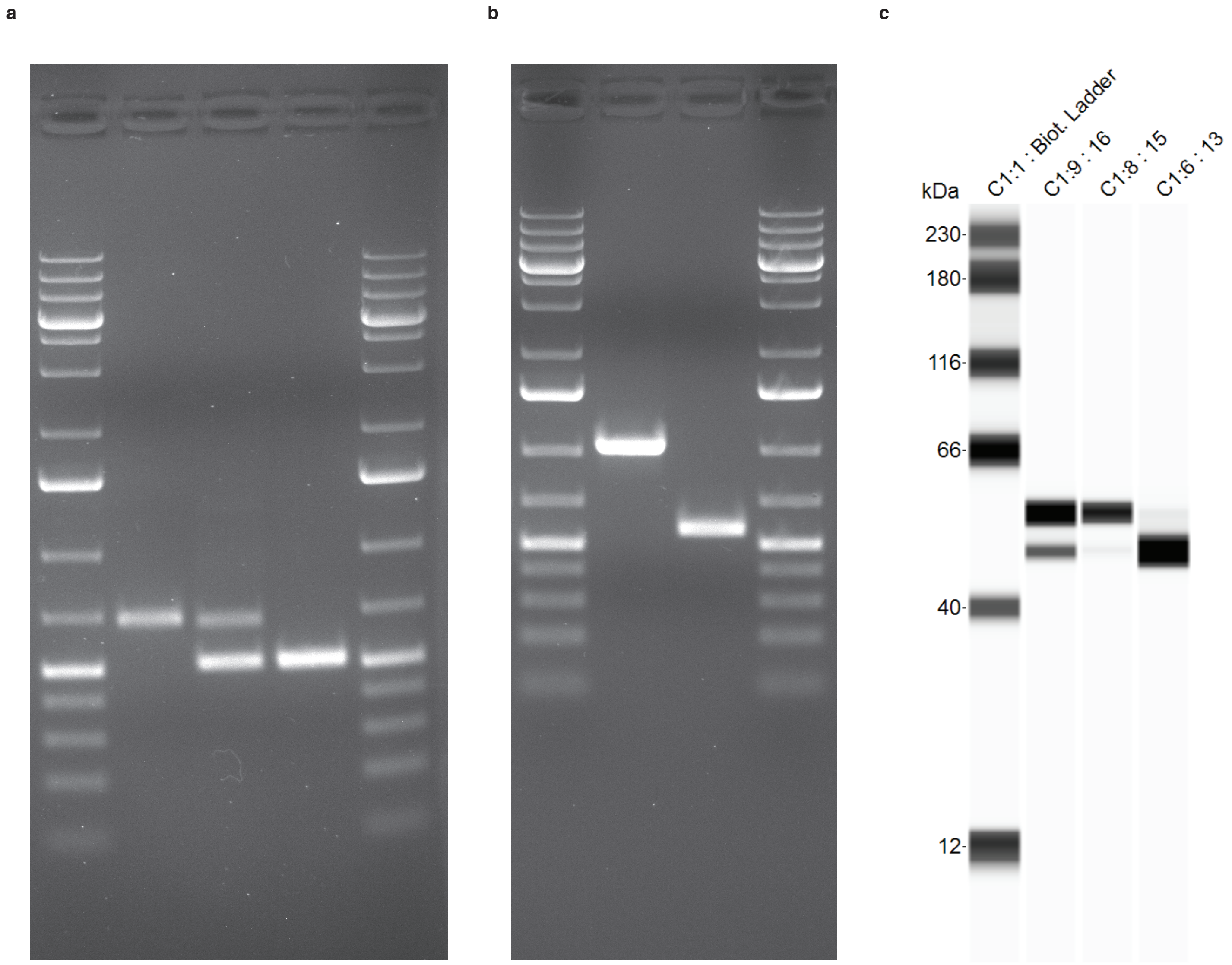
